# Characterization of expression elements for an AAV delivered antibody in nonhuman primates when co-delivered with PD-L1

**DOI:** 10.64898/2026.05.29.728808

**Authors:** Isai Leguizamo, Abubakarr A. Koroma, Michael Kuipa, Natalie S. Correa, Yash Barot, Sarah D. Hernandez, Meher Sethi, Anushka Das, Jun Xie, Guangping Gao, Stacey Weissman, Casey Whitehead, Stephanie Ehnert, Jennifer S. Wood, Priya Dhole, Matthew R. Gardner

**Affiliations:** Division of Microbiology and Immunology, Emory National Primate Research Center, Emory University, Atlanta, GA 30329 USA; Horae Gene Therapy Center, University of Massachusetts Medical School, Worcester, MA USA; Department of Genetics & Cellular Medicine, University of Massachusetts Chan Medical School, Worcester, MA, USA; Division of Animal Resources, Emory National Primate Research Center, Emory University, Atlanta, Georgia USA; Department of Medicine, Division of Infectious Diseases, Emory University, Atlanta, GA USA

## Abstract

Successful AAV-expressed antibody therapy for HIV-1 requires broadly neutralizing antibody (bNAbs) concentrations and reduced immune responses to sustain viral suppression without ART. We have previously demonstrated that co-delivery of AAV-expressed PD-L1 reduces immune responses against HIV-1 bNAbs in rhesus macaques. Here we systematically evaluated six AAV9 transgene cassettes encoding 10-1074 with different promoter/intron combinations (CMV, CMV/R, CBA, CASI, CB7, EF1α) across in vitro systems, immune-deficient mice, and in rhesus macaques. We show that both promoter and species selection, leads to differences in 10-1074 concentrations with the CB7 promoter leading to greatest expression in mice and CMV/R promoter in macaques. In addition to differences observed, loss of 10-1074 serum concentrations in macaques resulted in higher anti-drug antibody responses and antigen specific IFN-y T cell responses were focused on the 10-1074 heavy-chain variable region. Furthermore, inclusion of the WPRE greatly impacted 10-1074 expression leading to higher concentrations in both mice and nonhuman primates. Lastly, circulating 10-1074 in macaques retained neutralizing activity against diverse HIV-1 pseudovirus isolates. Together these results demonstrate how expression elements influence AAV-expressed antibodies in the context of co-delivery and highlight the need for further improvements to AAV transgene cassettes when co-delivered with AAV expressed PD-L1.

## INTRODUCTION

Adeno-associated virus (AAV)-delivered antibodies have emerged as a promising strategy for providing long-term protection against various infectious diseases. Unlike traditional vaccines that rely on eliciting endogenous immune responses for protection, AAV enables direct delivery of antibody encoding genes to host tissues such as skeletal muscle, resulting in sustained antibody expression from a single administered dose. Numerous studies have utilized this approach for protection against a wide array of viral pathogens, highlighting its versatility and therapeutic potential.^1–4^ Among these applications, using AAV for HIV-1 has gained particular traction because it allows for continuous expression of potent broadly neutralizing antibodies (bNAbs) potentially lasting several years.^5,6^ Multiple studies have demonstrated that bNAbs can suppress plasma viremia, delay viral rebound post-treatment interruption, and protect against infection.^7–13^ In one notable study, Martinez-Navio and Fuchs et al. demonstrated that a macaque achieved durable expression of an anti-SIV antibody following AAV1 administration for nearly seven years, maintaining concentrations of up to 200 μg/mL.^14^ Additionally, a recently completed Phase 1 clinical trial demonstrated the safety and feasibility of AAV8-delivered VRC07 in people living with HIV, with sustained expression lasting up to three years in some participants following a single administered dose.^15^

Despite early results highlighting the therapeutic potential of sustained antibody expression using AAV vectors, bNAbs often exhibit high levels of somatic hypermutation.^16–20^ While very critical for breadth and potency, these highly mutated variable heavy and light chain sequences increase the potential for developing host immune responses against the expressed antibody and transduced cell resulting in an impact to bNAb expression.^14,21–23^ In a previous study, we demonstrated that immune responses such as anti-drug antibodies (ADA) limit long-term expression of AAV-expressed HIV-1 bNAbs in a rhesus macaque model.^24^ To overcome this barrier, we recently employed the programmed cell death protein 1 and programmed cell death ligand 1 (PD-1/PD-L) immune checkpoint pathway, an interaction used by T cells to promote immune tolerance, to reduce immune clearance of AAV-transduced cells and the expressed antibody.^25–27^ Our recent study demonstrated that co-delivering AAV9 vectors expressing PD-L1 and AAV9 vectors encoding HIV-1 bNAbs (10-1074 or 3BNC117) overcame host immune responses, achieved durable prolonged expression of bNAbs at therapeutic concentrations, and provided protection against multiple SHIV_AD8-EO_ challenges in rhesus macaques.

While immune modulation using PD-L1 successfully enhanced the durability of bNAb expression, its expression is also heavily influenced by transgene elements such as promoters in the cassette itself.^28–31^ It’s known that promoter selection plays a critical role in influencing gene expression and kinetics, however no studies have compared these regulatory elements in the context of co-expression of both PD-L1 and an HIV-1 bNAb. In our previous study, 10-1074 expression was driven by the chicken beta actin (CBA) core promoter and PD-L1 was expressed using the cytomegalovirus (CMV) core promoter. The rationale for this approach was based on previous observations showing faster expression kinetics mediated by the CMV promoter compared to the CBA promoter.^32,33^ Thus, enabling earlier surface expression of PD-L1 in transduced cells and forming an immunological shield prior to 10-1074 secretion.

To further build on our previous work, the goal of this study was to further characterize AAV transgene cassettes for 10-1074 expression by systematically evaluating how core promoters impact 10-1074 expression. In addition, we explored the necessity of the woodchuck hepatitis virus post-transcriptional regulatory element (WPRE), which has been commonly used in AAV transgene cassettes to enhance mRNA stability and transgene expression.^34–36^ While the WPRE has been shown to enhance transgene expression, its necessity remains unclear, specifically when paired with transgene cassettes containing strong promoters. By evaluating both core promoters and the inclusion of the WPRE, we aimed to define how these transgene cassette elements impact 10-1074 expression, durability, and immunogenicity.

## RESULTS

### Characterization of 10-1074 expressed from transgene cassettes utilizing different promoter combinations

To evaluate the impact of core promoter selection on the expression of 10-1074, we initially designed five AAV9 transgene cassette plasmids containing distinct expression elements (**Fig. 1A**). Each cassette encoded the 10-1074 heavy and light chain separated by a P2A self-cleaving peptide and terminated with an SV40 late polyadenylation (polyA) signal. The P2A sequence was selected based on prior studies demonstrating improved expression efficiency compared to T2A or F2A peptides when used to link immunoglobulin heavy and light chains in AAV expressed antibody constructs.^21^ Additionally, consistent with our previous studies, we utilized the SV40 PolyA signal sequence for termination and the CMV enhancer each promoter. Unlike our previous studies, we removed the WPRE after the stop codon and prior to the SV40 PolyA signal sequence due to the size of CB7 promoter not being able to fit the WPRE within the packaging constraints of an AAV expression cassette. All cassettes were flanked by AAV2 inverted terminal repeats. The differences within the five expression cassettes were focused on the promoter and intron combinations. The CMV construct incorporated the CMV promoter (CMVp) followed by an SV40 intron (SV40i). The modified CMV/R construct contained an HLTV-1 R enhancer element and CMV intron (CMVi) downstream of the promoter region. The CBA, CASI, and CB7 designs utilized the CBA promoter in combination with either an SV40 intron (CBA), a synthetic ubiquitin C enhancer intron (CASI), or a chimeric intron (CB7). Lastly, the EF1α (human elongation factor 1 alpha) design employed its native EF1α promoter and EF1α intron. Vector genomes sizes ranged from 3.6 kb to 4.4 kb, sizing well within the AAV genome capacity. To assess whether our promoter/intron selections impacted in vitro neutralizing activity of 10-1074, we transiently transfected HEK293T cells with each AAV transfer plasmid or a double transfection of 10-1074 heavy and light chain expression plasmids. Neutralizing activity was determined against two HIV-1 pseudoviruses representing clade B (PVO.4) and clade C (CE1176) using TZM-bl neutralization assay (**Fig. 1B**). The native 10-1074, which served as a control and is denoted by the black curve, demonstrated potent neutralization activity against both pseudoviruses. As anticipated, all purified 10-1074 antibodies expressed from the five promoter AAV cassettes exhibited nearly overlapping neutralization curves with the control, demonstrating no loss of functional neutralizing activity throughout the different cassette designs. These results demonstrated that the differences in promoter and introns did not compromise the functional integrity of 10-1074 and are therefore suitable for subsequent in vivo studies.

**Figure 1.**
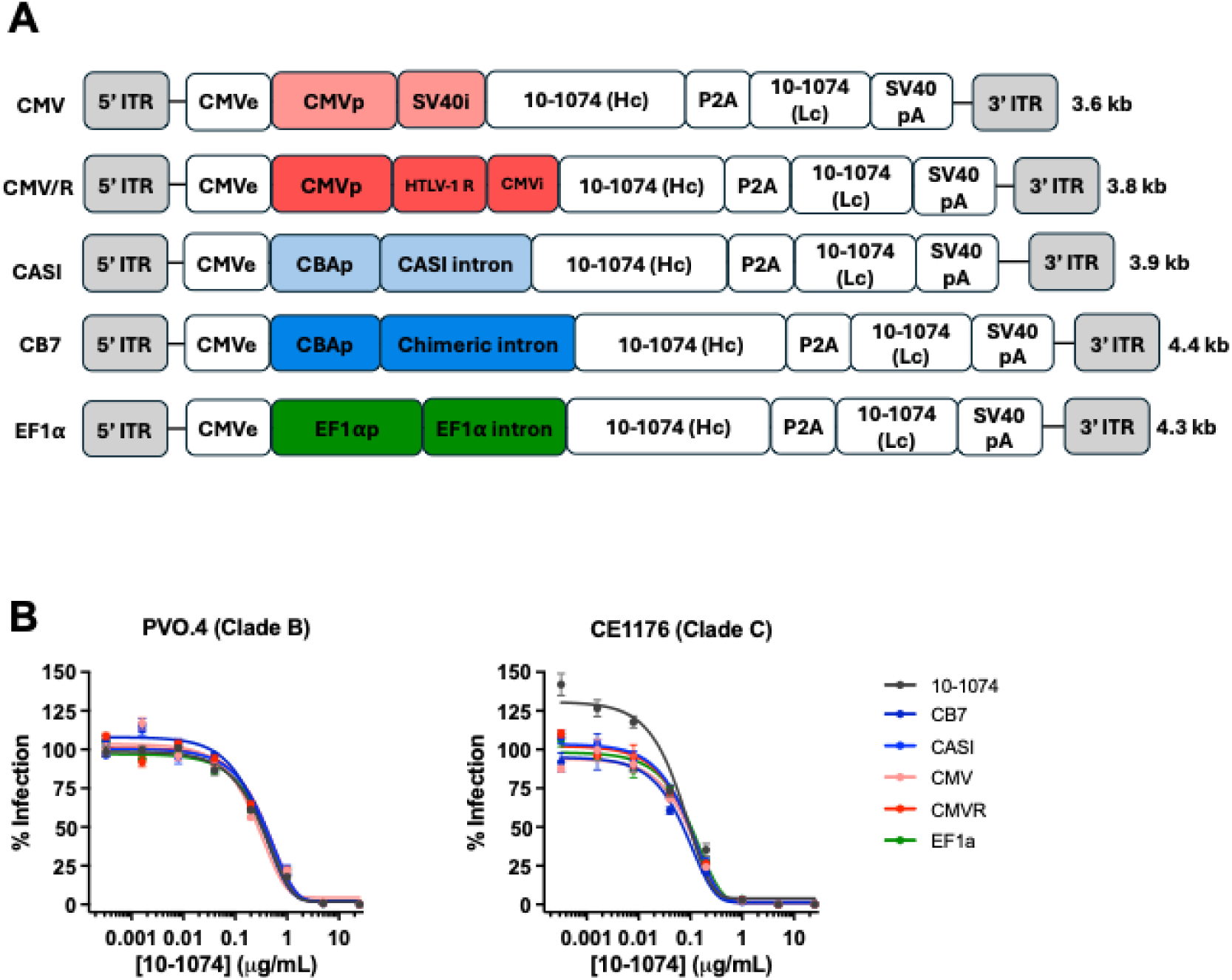
In vitro characterization of AAV9 transgene cassettes expressing 10-1074. (**A**) Schematic of the AAV9 transfer plasmids encoding 10-1074 heavy (Hc) and light chain (Lc). Constructs include CMV, CMV/R, CASI, CB7, or EF1α promoters (p) with corresponding introns (i). All transgene cassettes were flanked by AAV2 ITRs. (**B**) In vitro neutralization of 10-1074 expressed from each construct against a Clade B (PVO.4) and Clade C (CE1176) HIV-1 pseudovirus. Neutralization potency was determined by the TZM-l neutralization assay. Briefly, pseduovirus was pre-incubated with titrated amounts of antibody and incubated at for 1 hr at 37°C. Following incubation, TZM-bl cells were diluted to 100,000 cells/mL, added to the virus/antibody mixture, and incubated for 44 hrs at 37°C. Neutralization was determined by reduction of luciferase signal compared to virus plus cells alone. Error bars indicate S.D.

### Evaluation of AAV9-expressed 10-1074 in NSG mice

To evaluate the impact of promoter selection on in vivo 10-1074 expression, we tested antibody expression through intramuscular administration of AAV9 in NSG (NOD-*scid* IL2Rgamma^null^) mice. Purified AAV9 vectors expressing 10-1074 from different transgene cassettes encoding the five described promoters from Fig. 1A were administered into the left gastrocnemius muscle at a single dose of 2.5×10^12^ vg/kg (n = 4 mice per group). Plasma concentrations from the five groups were tracked over an 8-week period post AAV9 administration by gp120 ELISA **(Fig. 2A**). The CB7 group exhibited the highest plasma concentrations across all four mice over 8-weeks, ranging from 222 to 248 μg/mL at week 8. A slower but more gradual increase in 10-1074 concentration was observed in CMV/R, EF1α, and CBA groups over the course of the study. The CMV group, which utilizes the SV40 intron, exhibited the lowest concentrations ranging from 13 to 35 ug/mL at week 8. Group averages (**Fig. 2B**) confirmed robust differences in 10-1074 plasma concentration between the different groups with the CB7 group yielding the highest concentration (231 μg/mL) at week 8 followed by the CMV/R (126 μg/mL), EF1α (109 μg/mL), CASI (100 μg/mL), and CMV (21 μg/mL) group. Area under the curve (AUC) analysis (**Fig. 2C**) further supported these findings showing statistically significant higher 10-1074 concentrations in the CB7 group and statistically significant lower 10-1074 concentrations in the CMV group throughout the 8-week study. Together, these data demonstrate in vivo expression of 10-1074 is greatly impacted by selection of upstream promoter and intron configuration in NSG mice. Given these results we down-selected three constructs (CB7, CMV/R, and EF1α) for subsequent evaluation in nonhuman primates.

**Figure 2.**
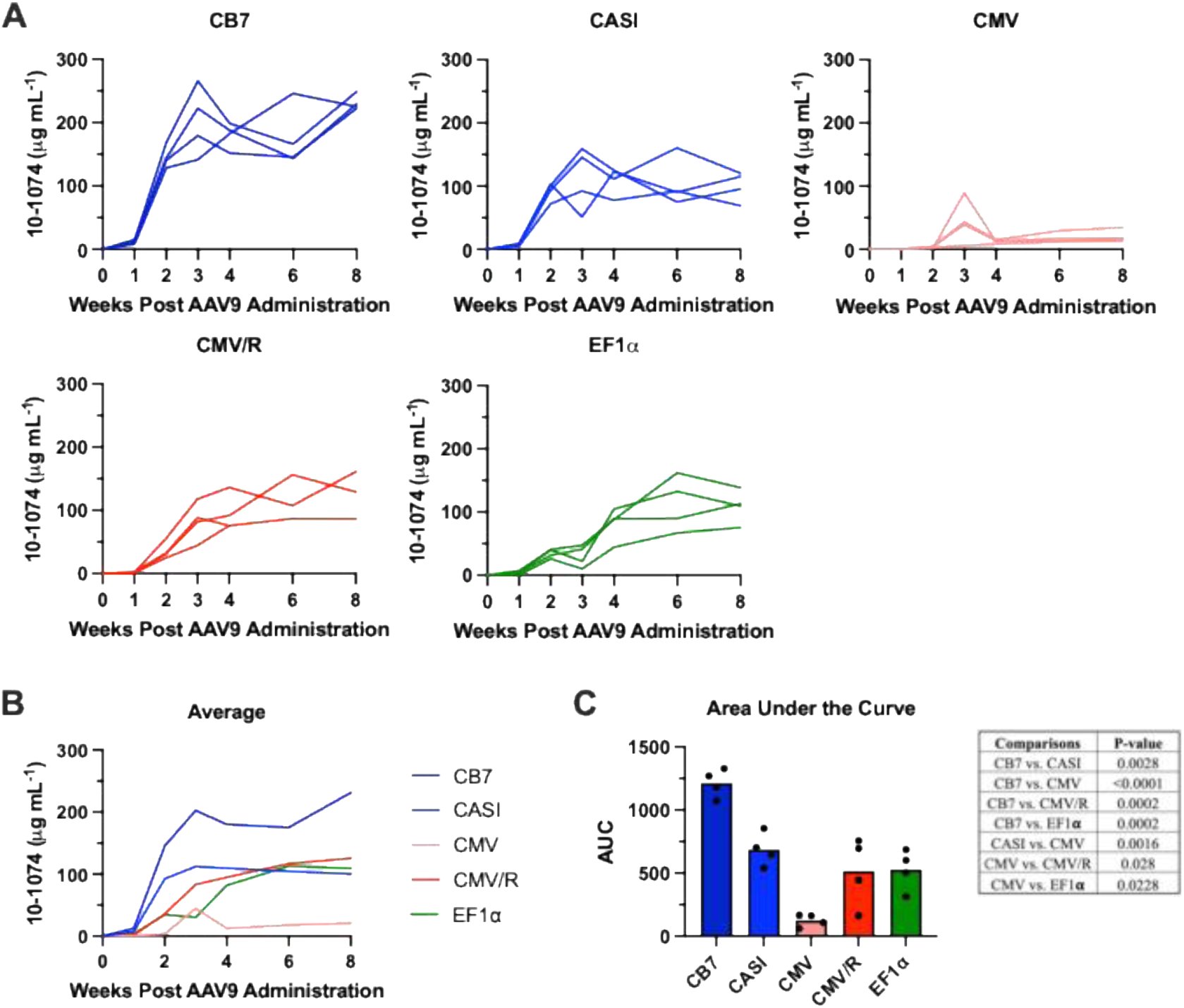
In vivo expression of 10-1074 from AAV9 vectors utilizing different promoters in NSG mice. (**A**) Serum concentrations of 10-1074 from five groups of NSG mice (n = 4) following a single intramuscular administration of AAV9 vectors at a dose of 2.5×10^12^ vg/kg. (**B**) Mean 10-1074 serum concentrations from all groups throughout eight weeks post AAV9 administration shown in (A). (**C**) Area under the curve (AUC) analysis of serum concentrations from each mouse per group. Each individual icon represents a mouse in that group. Statistical comparison of the promoter groups was determined by a one-way ANOVA with Tukey’s multiple comparison test. Statistically significant p-values are indicated in the Table.

### Evaluation of 10-1074 expression in NHPs following co-delivery with AAV9-PD-L1

AAV-delivered bNAbs in nonhuman primates has been plagued by host immune responses against the expressed bNAb. However, we have previously demonstrated that co-delivery of AAV9 expressed PD-L1 (AV9.PD-L1) can decrease host immunogenicity against 10-1074 in rhesus macaques. Our previous study utilized a dose of 2.5×10^12^ vg/kg for each AAV9 vector. Because of the results from the NSG mouse study, we reduced the dose of each vector (AAV9.PD-L1 and AAV9.10-1074) to 1.25×10^12^ vg/kg hypothesizing that we could still achieve the same concentrations with stronger promoters. We had three groups of four rhesus macaques each receiving their respective co-delivered AAV9 vectors (CB7, CMV/R, EF1α) expressing 10-1074 (AAV9.10-1074) and PD-L1 (AAV9.PD-L1) which utilizes the CMV promoter and SV40 intron. The vectors were administered evenly over eight IM injection sites: two per quadricep muscle, one per deltoid muscle, and one per bicep muscle. 10-1074 serum concentrations were quantified from serum samples taken over the course of 30 weeks by gp120 ELISA. Macaque weight ranged from 3.9 to 9.17 kg at the time of AAV9 administration and no significant weight loss was observed in any group (**Fig. S1**). As expected, following co-administration of AAV9.10-1074 (CB7, CMV/R, or EF1α) and AAV9.PD-L1, 10-1074 concentrations varied across groups (**Fig. 3A**). Surprisingly, in contrast from the previous observation in NSG mice, the CB7 group yielded the lowest 10-1074 serum concentrations overall, with an average range of 9 to 19 μg/mL throughout the 30-week study. One macaque, Mm038, had a peak concentration of 18 μg/mL at week 3 before dropping to below limits of detection by week 10. The CMV/R group achieved the highest 10-1074 serum concentrations with three of four animals sustaining 24, 43, and 93 μg/mL at week 30 post administration. Again, one macaque, Mm038 had a week 4 peak concentration of 31 μg/mL before dropping to below the limits of detection at week 6. Similar to the CMV/R, we observed more variable 10-1074 concentrations across the EF1α group. Like the other two groups, one macaque, Mm040, peaked at week 6 with 10-1074 concentrations reaching 133 μg/mL before falling to below the limits of detection at week 6. Mm042 peaked at week 6 (75 μg/mL) and gradually decreased to 12 μg/mL by week 30. Mm041 peaked at week 10 (69 μg/mL) and decreased to 36 ug/mL by week 30. Interestingly, Mm039, peaked at week 14 (162 μg/mL) before gradually dropping to 73 μg/mL. Group average 10-1074 serum concentrations and AUC analysis (**Fig. 3B, C**) further demonstrated that the CMV/R group achieved the highest concentration over time followed by the EF1α and CBA groups. To evaluate whether host variables contributed to the observed differences seen in macaques, we correlated serum 10-1074 AUC with sex, age, and body weight and found no associated significance (**Fig. S2**). Lastly, to compare promoter performance between species, we evaluated the AUC of 10-1074 serum concentrations between NSG mice and rhesus macaques from weeks one to eight (**Fig. 3D**). The CMV/R and EF1α promoters exhibited relatively consistent 10-1074 expression activity across both species, with comparable AUCs in mice and rhesus macaques. In contrast, the CB7 promoter, which drove the highest 10-1074 expression in NSG mice, showed a lower AUC, indicating a significant loss in promoter activity throughout the study. These findings demonstrated that in addition to promoter selection, different species lead to differences in expression of 10-1074.

**Figure 3.**
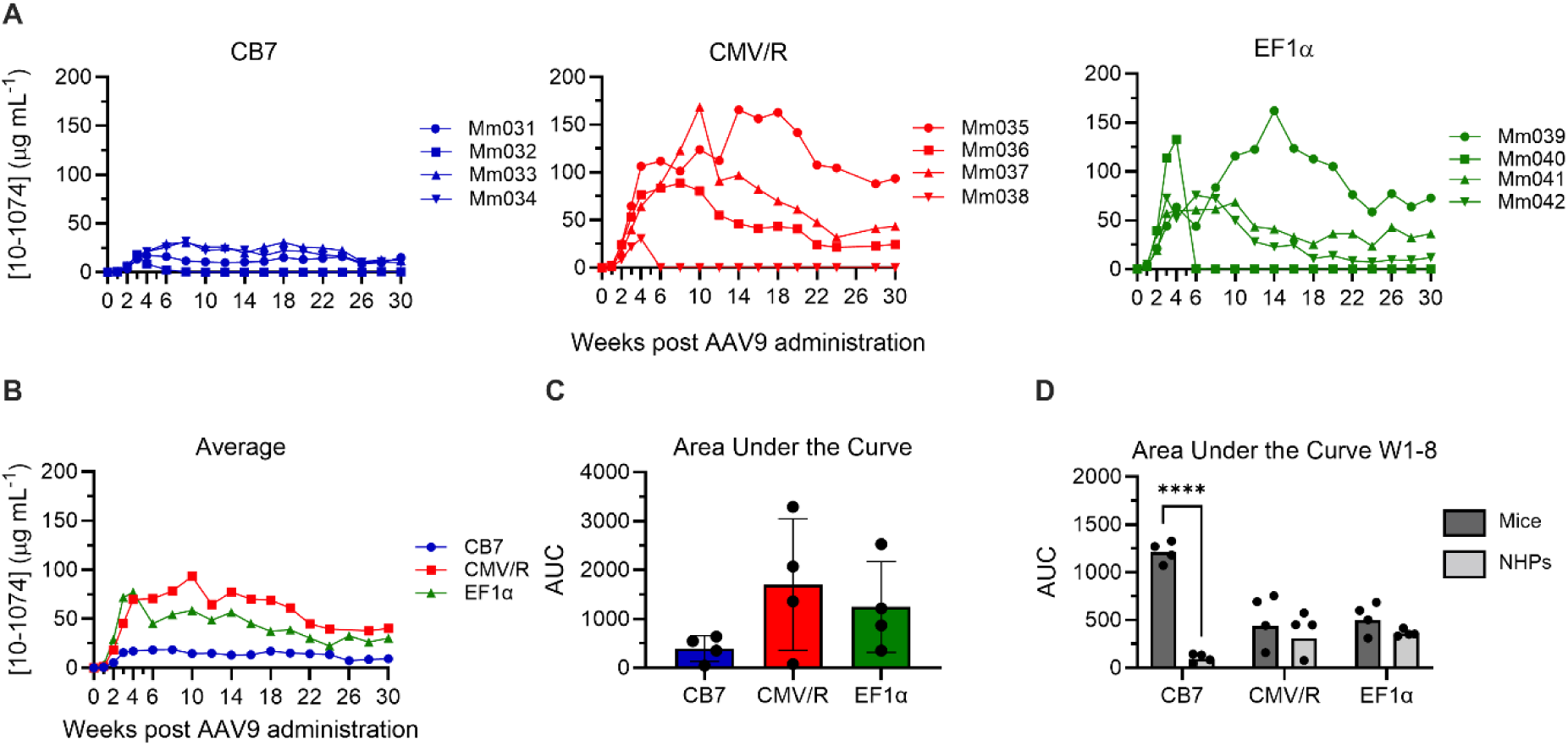
Comparison of 10-1074 expression rhesus macaques following AAV9 administration comparing down-selected promoter cassettes. (**A**) Serum 10-1074 concentrations from 3 groups of rhesus macaques (n = 4) following co-delivery of indicated AAV9.10-1074 and AAV9.PD-L1 (CMV promoter) at a dose of 1.25×10^12^ vg/kg per vector. (**B**) Group averages of 10-1074 serum concentrations from all groups throughout the study. (**C**) Average AUC analysis of serum concentrations from each group over 30 weeks. Each dot represents one macaque. (**D**) Comparative average AUC from weeks 1-8 in NSG mice (Fig. 2) versus rhesus macaques. Each dot represents a single animal. Statistical comparison of the promoter groups between species was determined by a two-way ANOVA with Tukey’s multiple comparison test. Significance defined as **** indicates a p-value ≤ 0.0001.

### Evaluation of host immune responses against AAV9-delivered 10-1074 and PD-L1 in NHPs

As previously mentioned, a major issue that continues in the field of AAV is the development of host immune responses such as anti-drug antibody (ADA) responses against expressed transgenes. In a previous study, we demonstrated that co-delivery of AAV-expressed PD-L1 overcame host immune responses against HIV-1 bNAbs in rhesus macaques. Given our previous results, we next analyzed whether ADA responses were generated against 10-1074 through the entire study **(Fig 4A)**. In this study, the CB7 group had one macaque (Mm032) that exhibited higher ADA responses shown as endpoint titers above the values determined at the time of AAV9 administration. However, these titers remained well below the highest measurable endpoint titer (1:1250). In contrast, one macaque in the CMV/R group (Mm038) developed a high ADA response that exceeded the highest endpoint titer at week 8. Similarly, one macaque in the EF1α group (Mm040) showed increasing ADA responses eventually peaking at the highest endpoint titer at week 18. Notably, all macaques with ADA responses above the highest endpoint titers corresponded to macaques that resulted 10-1074 serum concentrations <1μg/ml (**Fig. S3**), suggesting that elevated ADA responses were associated with loss of transgene expression. Interestingly, the macaque (Mm032) from the CB7 group that had low serum concentrations of 10-1074 did not develop ADA responses above the highest endpoint titer, supporting the idea that low antigen exposure may not be sufficient for ADA generation. In addition to ADA responses, we have previously shown the infiltration of T cells into the sites of AAV administration. Given the role of T cells driving antibody responses and killing transduced cells, we next evaluated the cellular immune responses against 10-1074 and PD-L1. We used an IFN-γ ELISpot assay by means of overlapping peptide pools spanning PD-L1, LS-Furin-P2A, and the 10-1074 heavy and light chains to determine if any regions contributed to T cell responses (**Fig. 4B**). Antigen-specific cellular immunity was observed against the 10-1074 heavy chain variable region (10-1074 Var H) at week 30, particularly in the CB7 group. The strongest IFN-γ response occurred in macaque Mm031, which also exhibited no measurable 10-1074 serum concentration and a steady rise in anti-10-1074 binding antibodies. Furthermore, responses to PD-L1, LS-P2A, and 10-1074 10-1074 Var L remained minimal across all groups. SEB elicited strong positive control responses, confirming T cell functionality.

**Figure 4.**
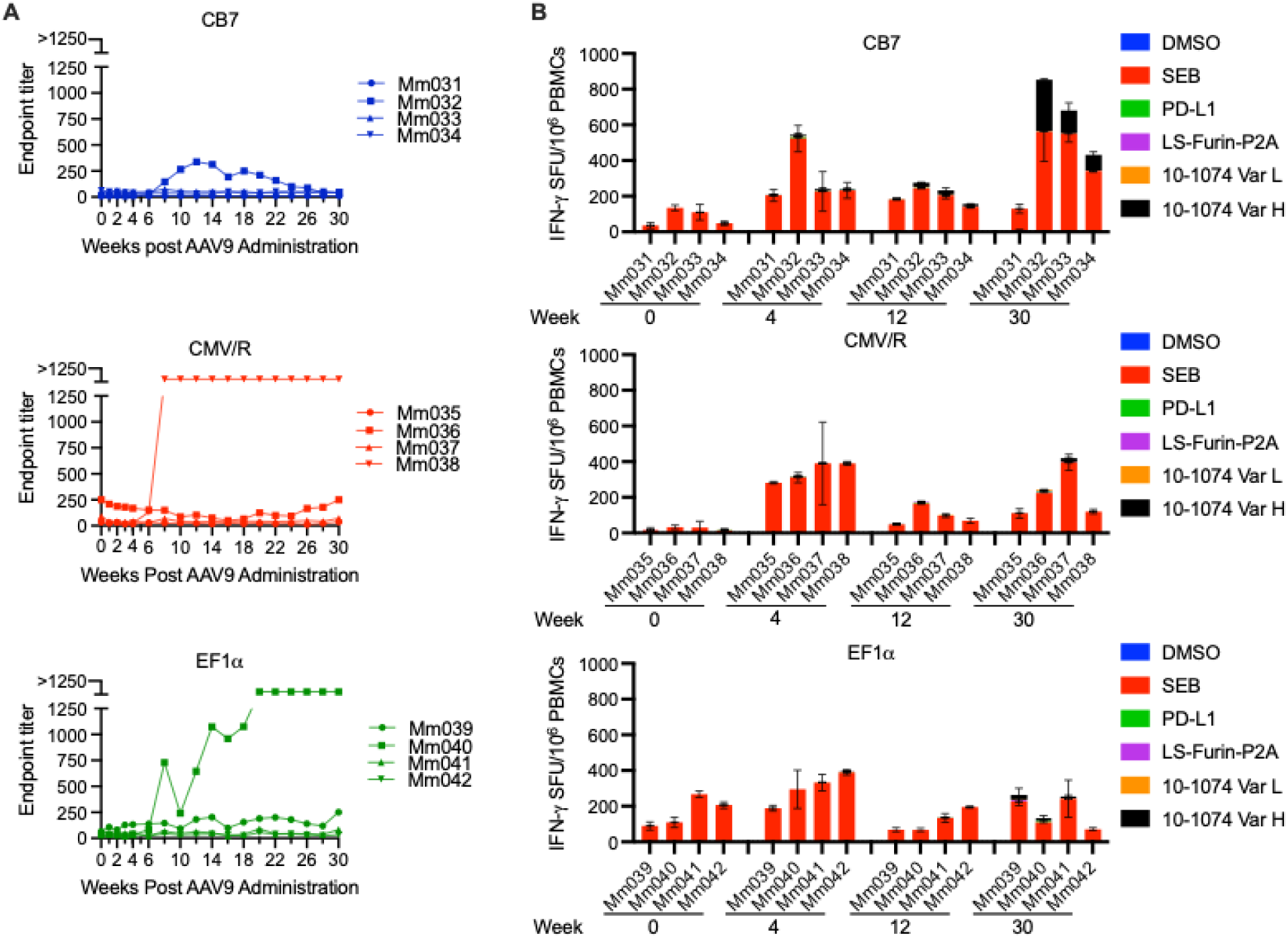
Host Anti-drug antibody and T cell responses against AAV9-delivered 10-1074 in rhesus macaques. (**A**) 30-week longitudinal ADA responses against 10-1074 in rhesus macaques receiving AAV9.10-1074 driven by the indicated promoter. Endpoint titers are defined as the serum dilution where OD_450_ = 0.2 as determined by ELISA. Macaques with ADA endpoint titers above the highest dilution measured were given a value of 1300. (**B**) IFN-γ ELISpot responses in macaques to indicated peptide pools derived from both PD-L1 and 10-1074 (LS-Furin-P2A, variable heavy chain, variable light chain) at weeks 0, 4, 12, and 30 following AAV9 administration. Staphylococcal enterotoxin B (SEB) was used at a positive control and DMSO as a negative control. Error bars indicate S.D.

### Evaluation of the WPRE for 10-1074 expression and host immune responses

Although the WPRE has been commonly used in AAV transgene cassettes to enhance mRNA stability and overall transgene expression, its necessity in the context of strong promoter designs and co-delivery with AAV9.PD-L1 remains unclear. Our previous study included the WPRE in the AAV9 10-1074 transgene cassette. To determine the necessity of the WPRE, we utilized the same cassette and AAV9 capsid used in our previous study but removed the WPRE (AAV9.CBA(- )WPRE.10-1074 **(Fig. 5A)**. We initially evaluated 10-1074 expression in NSG mice by intramuscularly administering the AAV9 vectors at a dose of 2.5×10^12^ vg/kg and tracking 10-1074 plasma concentrations over an 8-week period by gp120 ELISA. The (+)WPRE group revealed 10-1074 concentrations ranging from 281 to 760 μg/mL at week 8 post AAV9 administration (**Fig. 5B**). Analysis of group averages (**Fig. 5C**) revealed higher 10-1074 plasma concentrations in the (+)WPRE group, reaching a 5-fold difference of expression by week 8. Additionally, AUC analysis (**Fig. 5D**) demonstrated that the (+)WPRE group had significantly greater promoter activity throughout the 8-week period.

**Figure 5.**
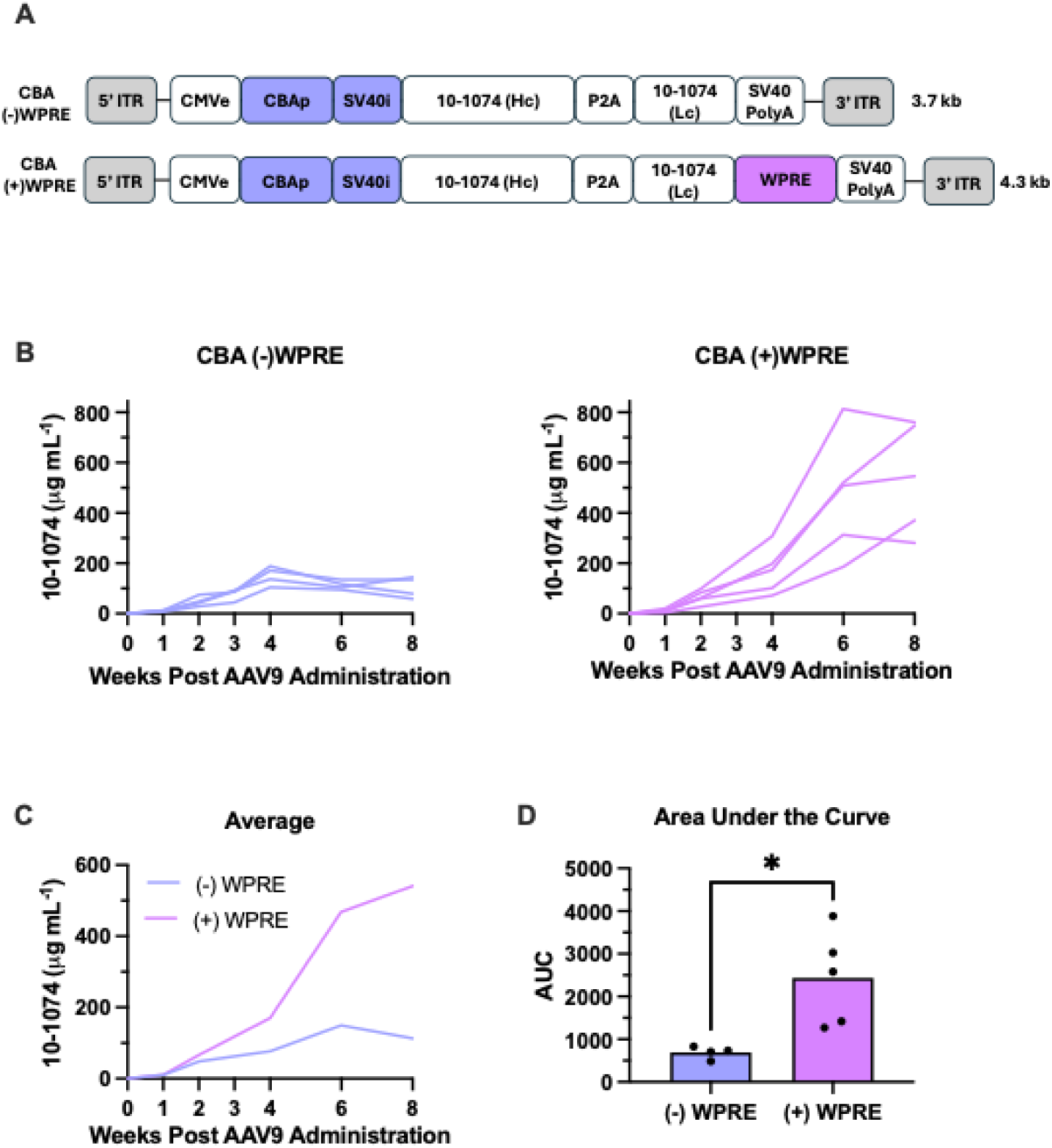
Characterization of 10-1074 transgene cassettes with or without the WPRE in NSG mice. (**A**) Schematic of CBA promoter-driven AAV9 transfer plasmids encoding 10-1074 containing or lacking the WPRE. (**B**) Serum concentrations of 10-1074 from two groups of NSG mice following a single intramuscular administration of AAV9.10-1074 with or without the WPRE at a dose of 2.5×10^12^ vg/kg. (**C**) Mean 10-1074 serum concentrations from both groups throughout eight weeks demonstrating greater expression with inclusion of the WPRE. (**D**) Average AUC analysis comparing 10-1074 concentrations throughout the 8-week study. Statistical comparisons between both groups were determined by an unpaired two-tailed t test with each dot representing AUC for an individual mouse. Significance defined as * indicates a p-value ≤ 0.05.

Given the species-specific differences we observed with the CB7 promoter, we sought to confirm the WPRE-specific differences observed in mice with rhesus macaques. Four rhesus macaques were administered AAV9.CBA(-)WPRE.10-1074 and AAV9.PD-L1 vectors at a dose of 2.5x10^12^ vg/kg of each vector. These conditions were the same as our previous study where the only difference was the inclusion of the WPRE in the AAV9.10-1074 vector cassette. In this group, all macaques achieved sustained expression of 10-1074 with serum concentrations ranging from 50 to 126 μg/mL by week 30 (**Fig. 6A**). However, our historical data demonstrated much higher serum concentrations ranging from 173 to 480 μg/mL by week 30 (**Fig. 6B**). Group averages and AUC analysis further demonstrated the differences in expression between both groups indicating that including the WPRE in our transgene cassette significantly enhances expression over time (**Fig. 6C**). AUC analysis comparison of (+) or (-) WPRE between species from weeks one to eight revealed significantly higher promoter activity in mice compared to macaques, however activity remained relatively consistent between NHPs (**Fig. 6D**). Interestingly, when assessing ADA responses in these groups, we observed no ADA responses above endpoint titers prior to AAV9 administration in the (-)WPRE group (**Fig. 7A**). In contrast, ADA endpoint titers were above baseline titers in the (+)WPRE group, prior to AAV9 administration and between weeks 10-20 post AAV9 administration, suggesting possible pre-existing immunity to AAV9. Nonetheless, these ADA titers in (+)WPRE group did not appear to impact 10-1074 expression. Consistent with our previous findings, IFN-γ responses were once again prominent against the 10-1074 variable heavy chain region (10-1074 Var H) at week 30. (**Fig. 7B).**

**Figure 6.**
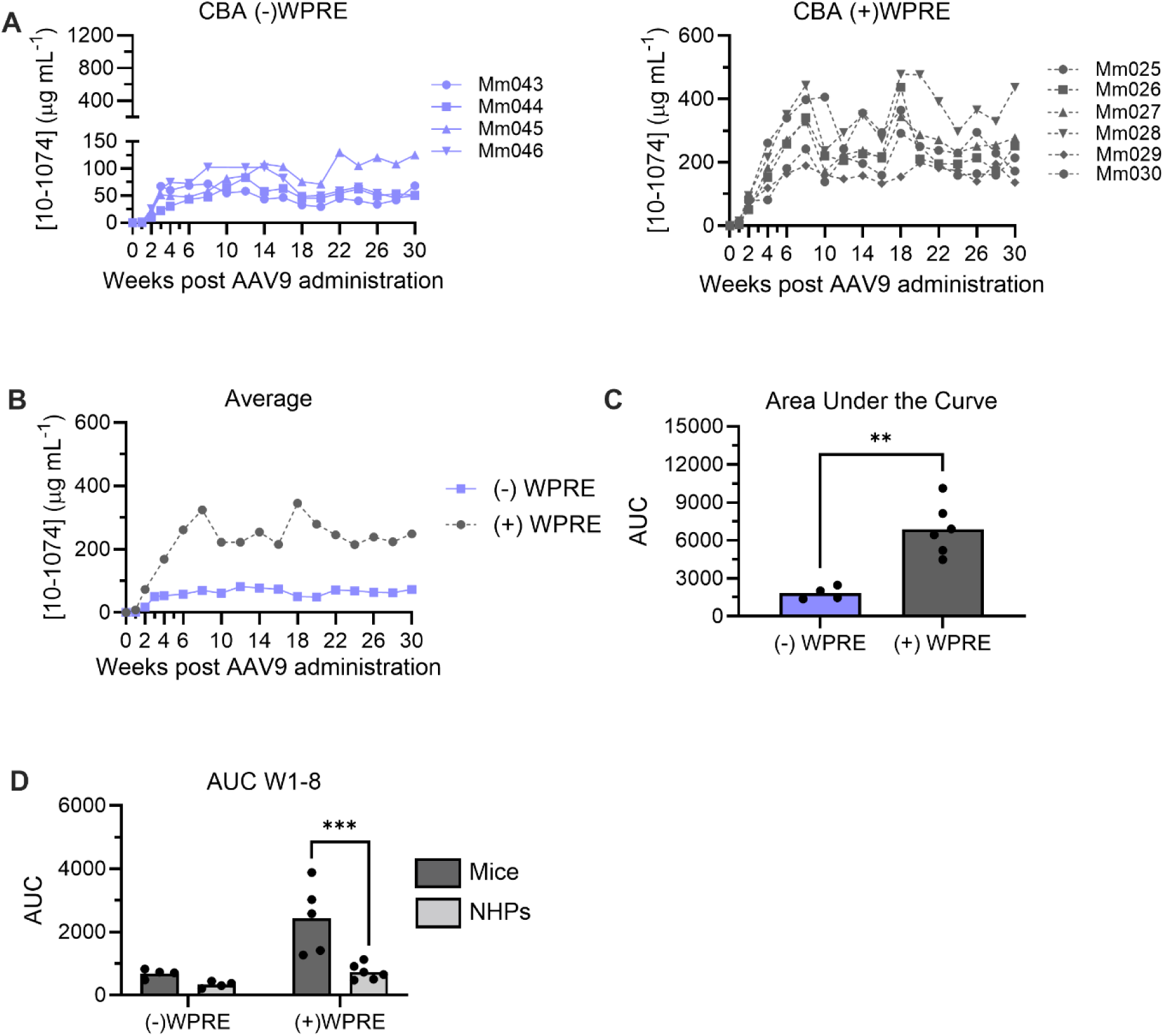
Comparison of CBA-driven 10-1074 expression with or without the WPRE in rhesus macaques. (**A**) Serum 10-1074 concentrations from rhesus macaques following co-delivery of AAV9.PD-L1 (CMV promoter) with AAV9.10-1074 (- WPRE, left) or AAV9.10-1074 (+ WPRE, right) at a dose of 2.5×10^12^ vg/kg per vector. Note that the +WPRE group (right) is based on historical data from Kuipa et al., 2026. (**B**) Group averages of 10-1074 serum concentrations from both groups throughout 30 weeks post AAV9 administration. (**C**) AUC analysis of serum concentrations. Statistical comparison was determined by an unpaired two-tailed t test. Significance defined as ** indicates a p-value ≤ 0.01. (**D**) Comparative AUC from weeks 1-8 in NSG mice versus rhesus macaques with or without the WPRE. Statistical comparison was determined by a two-way ANOVA with Tukey’s multiple comparison test. Statistical significance defined as *** indicates p-value ≤ 0.001).

**Figure 7.**
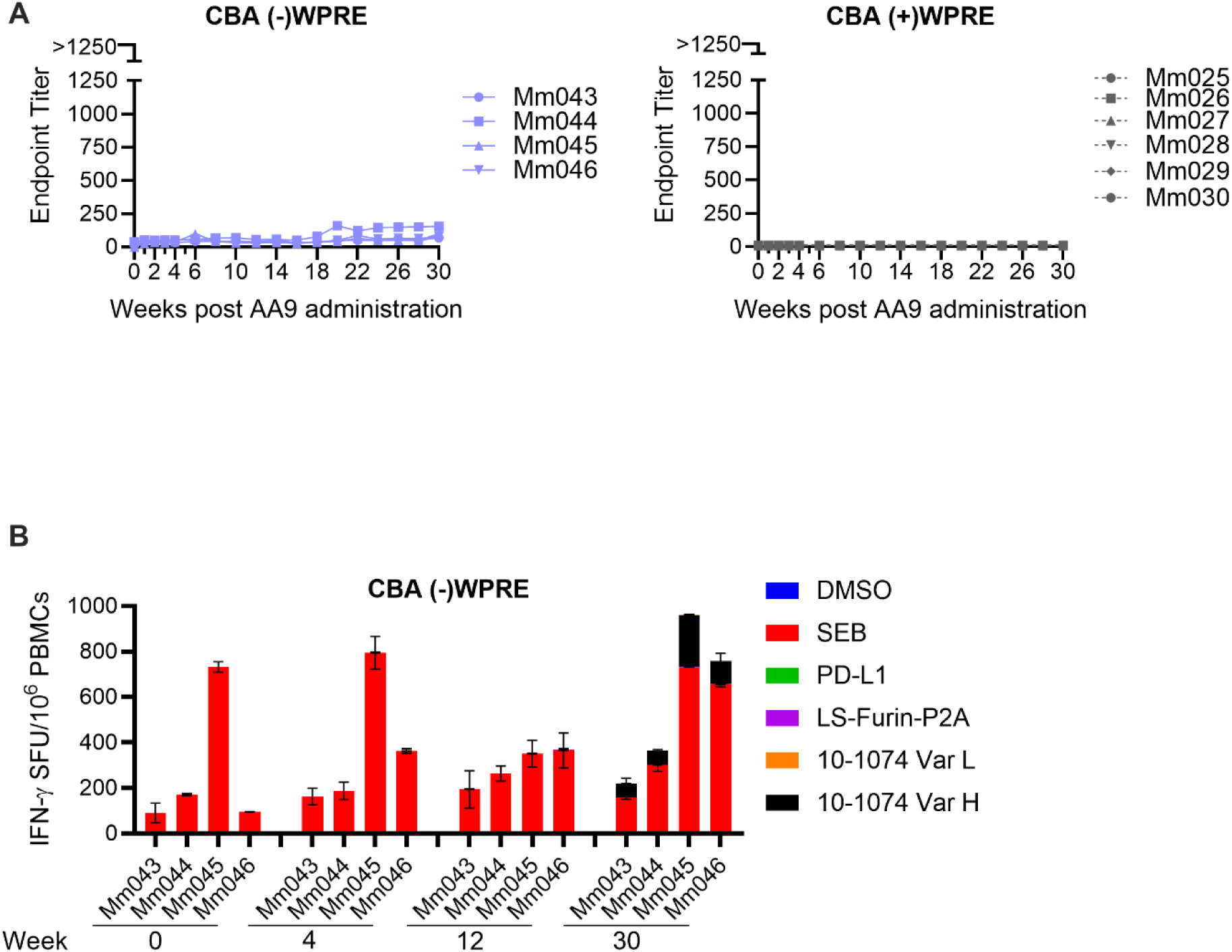
Characterization of host immune responses against expressed 10-1074 without the WPRE. (**A**) Longitudinal 30-week ADA responses in macaques receiving AAV9.10-1074 with or lacking the WPRE. Endpoint titers are defined as the serum dilution where OD_450_ = 0.2 as determined by ELISA. Note that the +WPRE group (right) is based on historical data from Kuipa et al., 2026. (**B**) IFN-γ ELISpot responses from macaques in the -WPRE group to peptide pools derived from both PD-L1 and 10-1074 (LS-Furin-P2A, variable heavy chain, variable light chain) at weeks 0, 4, 12, and 30 following AAV9 administration. Staphylococcal enterotoxin B was used at a positive control and DMSO as a negative control.

### Assessing neutralizing activity of 10-1074 and AAV in macaque serum

Serum samples from all macaques in these studies were analyzed to ensure that neutralizing activity was maintained ex vivo. Sera from the week 30 timepoint was initially diluted 1:10 in media then serially diluted 5-fold and mixed with either Clade A (BG505-T332N), Clade B (PVO.4), and Clade C (CE1176) HIV-1 pseudoviruses. SIVmac239 and pre-sera was used as a negative control. All macaques with measurable 10-1074 serum concentrations demonstrated neutralizing activity across all HIV-1 pseudoviruses (**Fig. 8**). Importantly, serum 10-1074 serum concentrations at week 30 correlated strongly with neutralizing activity across all three pseudoviruses. (**Fig. S4**). Additionally, as expected, we observed no neutralizing activity across all groups against SIVmac239, indicating no non-specific antiviral activity in macaque serum and confirming the specificity of 10-1074 to the V3-glycan supersite of HIV-1.

**Figure 8.**
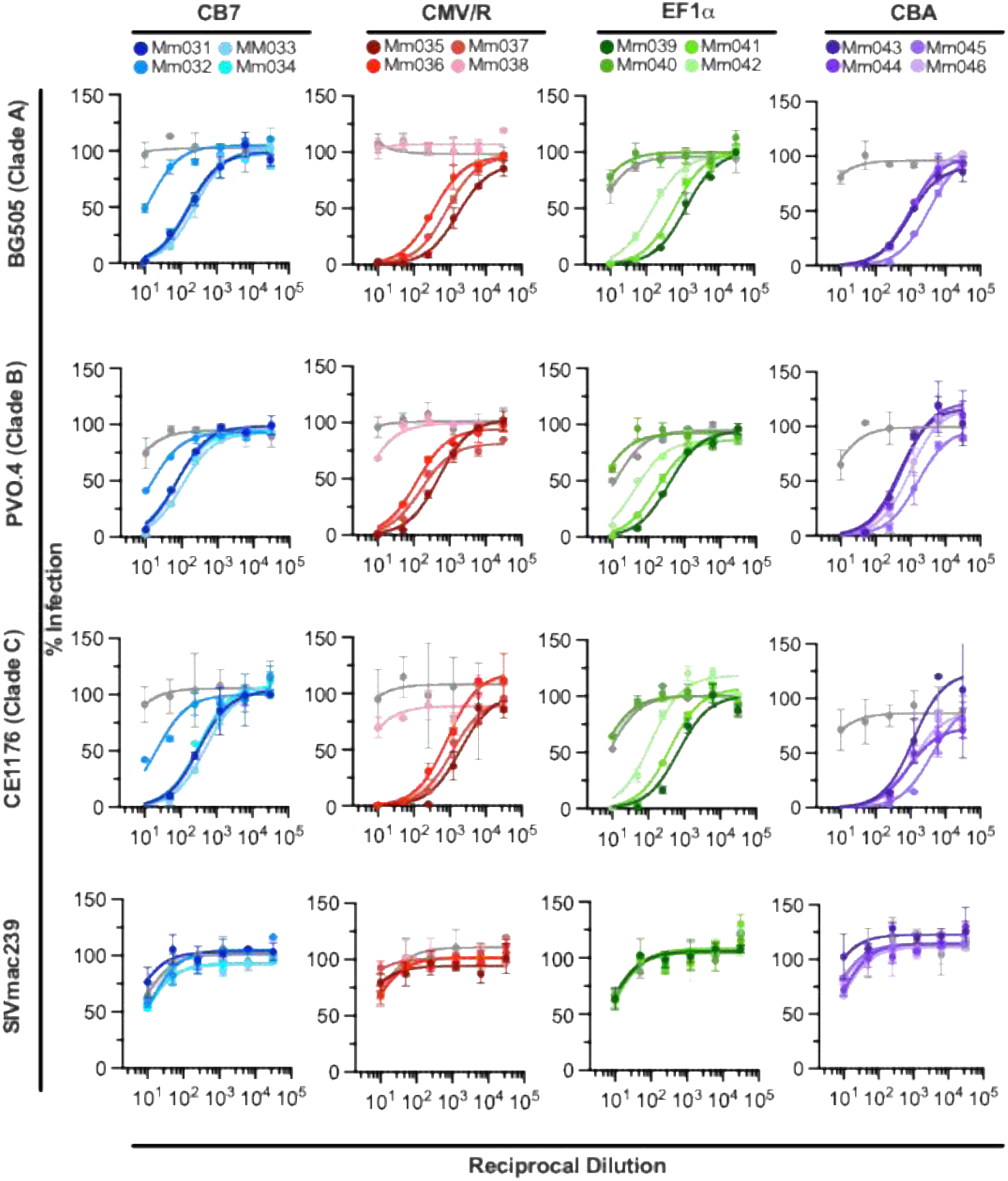
Neutralizing activity of AAV9-expressed 10-1074 in macaque serum against diverse HIV-1 strains. Ex vivo neutralizing activity from serum samples at 30 weeks post-AAV9 administration were performed against clade A (BG505-T332N), clade B (PVO.4), and clade C (CE1176) pseudoviruses by TZM-bl neutralization assay. A serum sample (pre-sera) from each group prior to AAV9 administration was used as a negative control. SIVmac239 pseudovirus was used to determine off-target luciferase reduction mediated by the serum sample to confirm the specificity of 10-1074 against the HIV-1 pseudoviruses. Error bars indicate S.D.

## DISCUSSION

Efficient and durable in vivo expression of HIV-1 bNAbs remains a major goal in the development of AAV gene therapy applications for both HIV prevention and cure strategies. Our previous study demonstrated that co-administration of AAV9 vectors expressing PD-L1 alongside AAV9-delivered bNAbs improved the durability of transgene expression through overcoming host immune responses. However, in this study our goal was to investigate how characterizing transgene cassettes, when co-delivered with AAV9 vectors expressing PD-L1, impact in vivo expression and immunogenicity of AAV9-delivered 10-1074. In our first approach, we selected various promoters/introns for expression of 10-1074. Our data confirms that promoter selection plays a critical role in 10-1074 expression, particularly in the context of different species. In NSG mice, we observed the highest 10-1074 concentrations in the CB7 group **(Fig. 2A)**, supporting previous reports highlighting CB7’s strength in muscle tissue. However, this promoter failed to translate to high expression in rhesus macaques, potentially suggesting species-specific differences in transcription factor availability. We observed an 11-fold decrease from mice to monkeys, like what was previously shown by Greig et al. who reported a 4-fold decrease from mice to monkeys in their study.^37^ In contrast, the EF1α and CMV/R promoters resulted in relatively consistent expression across both species, with the CMV/R achieving the highest sustained 10-1074 serum concentrations in macaques. The findings underscore the importance of validating promoters in relevant animal models, as promoter strength may vary among species.

An important consideration in our study is the impact of promoter expression kinetics between our PD-L1 and 10-1074 AAV cassettes. In our previous study, we utilized the CMV promoter for PD-L1 and CBA promoter for 10-1074 expression. This selection was based on previous observations that CMV exhibits faster kinetics of expression than CBA, resulting in earlier surface expression of PD-L, inhibiting PD-1 signaling, and providing early protection prior to secretion of 10-1074. However, in this study, the promoters driving 10-1074 expression varied in strength relative to the CMV promoter used for PD-L1 expression. For example, previous reports show CB7, EF1α, and CMV/R promoters are stronger than CMV. These differences in promoter potency may have resulted in earlier expression of 10-1074, preceding surface expression of PD-L1 and reducing the intended immunological “shielding” effect of AAV9-transduced muscle cells. Additionally, co-delivery of two AAV vectors with strong promoters raises the possibility of promoter competition for transcriptional machinery which may further influence overall transgene expression. Because differences in promoter strength and expression kinetics can influence the magnitude and timing of antigen exposure, these factors may have also played a role in the development of ADA and T cell responses in our groups. Our findings further supported that loss of bNAb expression is attributed to higher ADA responses. This was most notable in one macaque (Mm038) from the CMV/R group and another (Mm040) in the EF1α group. It’s also possible that absent or low antigen exposure may potentially limit ADA formation as seen with one macaque (Mm032) in the CB7 group.

The WPRE has been commonly used in AAV transgene cassettes to enhance mRNA stability and overall transgene expression. However, numerous published studies have conflicting findings on whether the WPRE is necessary. Thus, our second approach aimed to determine the necessity of the WPRE for 10-1074 expression when co-delivered with AAV.PD-L1 in rhesus macaques via IM administration. Our data revealed that the WPRE significantly improved 10-1074 expression in both mice and rhesus macaques. Notably, in the CBA group lacking the WPRE, no macaques experienced a loss of 10-1074 expression to serum concentrations less than 1 μg/mL. In fact, all macaques achieved concentrations of therapeutic magnitude that have been shown to suppress viremia when combined with other bNAbs. Interestingly, when compared to our historical data, we observed a trend of higher ADA responses, specifically between weeks 10-20 post AAV9 administration, compared to the CBA group lacking the WPRE. However, in both groups, ADA responses did not exceed our highest endpoint titer. These results demonstrate that although the inclusion of the WPRE enhanced antigen expression and potentially increased immune recognition, it did not drive ADA responses sufficient to cause rapid clearance of 10-1074. Together, these results further support the incorporation of the WPRE into bNAb-expressing transgene cassettes for future in vivo studies.

While quantifying 10-1074 serum concentrations and evaluating host immune responses gave us insights into the durability and immunogenicity of our AAV-expressed bNAb, it was equally important to confirm that circulating 10-1074 retained its functional activity. Our results demonstrated that 10-1074 retained its neutralizing activity throughout the 30-week study and that greater expression of 10-1074 correlated with greater neutralizing activity. Our findings also confirmed the in vivo breadth of 10-1074 against Clade A (BG505-T332N), Clade B (PVO.4), and Clade C (CE1176) pseudoviruses, an objective yet to be obtained by conventional vaccine designs.

Overcoming the host immune response to achieve prolonged and high in vivo expression of bNAbs has been a major hurdle in the field of AAV gene therapy applications for HIV. However, our previous study was the first to demonstrate that co-administration of AAV9 vectors expressing PD-L1 alongside AAV9-delivered bNAbs significantly enhanced antibody expression through overcoming host immune responses. This present study was the first to investigate how different transgene cassettes, when co-delivered with AAV9 vectors expressing PD-L1, impact in vivo expression and immunogenicity of AAV9-delivered 10-1074. We demonstrated the importance of considering promoter strength when co-delivering two AAV transgene cassettes and additional factors that may influence transgene expression. Given our results, future studies may focus on engineering cassettes all, containing the WPRE, with a stronger promoter on PD-L1 and determine if transgene cassettes containing weaker promoters are able to achieve high and prolonged expression of bNAbs. We believe this study is a big step forward in the field of AAV-expressed antibodies and will serve as a framework for subsequent work.

## Materials and Methods

### Rhesus macaques

Sixteen Indian-origin rhesus macaques (10 males, 6 females), aged 2.75–6.2 years at the time of AAV administration, were enrolled in this study. Macaques were housed at the Emory National Primate Research Center (ENPRC) in Atlanta, Georgia. All protocols were approved by the Emory University Institutional Animal Care and Use Committee (Permit number: 202100453) and were performed in accordance with guidelines established by the United States Department of Agriculture (USDA) Animal Welfare Act and the recommendations of the National Institutes of Health (NIH) Guide for the Care and Use of Laboratory Animals (8th Edition). The animal care facilities at ENPRC are accredited by both the USDA and AAALAC, International. Macaques were housed in pairs with compatible animals for the entirety of the study. Macaques were screened for serum neutralizing antibodies against the AAV9 serotype prior to study initiation. Body weights at the time of vector administration ranged from 3.92–8.92 kg.

### Cell lines and plasmids

HEK293T cells were obtained from ATCC and grown in DMEM with 10% fetal bovine serum at 37°C. TZM-bl cells were obtained through the NIH AIDS Reagent Program, Division of AIDS, NIAID, NIH from Dr. John C. Kappes, Dr. Xiayun Wu, and Tranzyme and grown in DMD with 10% FBS at 37°C. AAV9.CAG.fLuc transfer plasmid was obtained from Addgene (#93281) and described previously. Rep/Cap expression plasmid for AAV9 was previously described. pHelper plasmid was from ATCC. The expression plasmids for 10-1074 heavy and light chains and AAV transgene cassette encoding rhesusized 10-1074 were previously described. For replacing the CMV promoter with CB7, CASI, CMV/R, and EF1a, each promoter was synthesized with the 5’ ITR CMV enhancer and respective intron by GenScript and inserted using the DraIII and PspOMI restriction sites into the previously described CBA construct. The 10-1074 transgene was cut out from the CBA cassette using NotI sites and inserted into cassettes containing AAV2 ITRs and the respective promoter CB7, CASI, and no WPRE. Additionally, the WPRE was removed from the original CBA 10-1074 cassette by synthesizing a fragment from the EcoRV cut site in 10-1074 to the MfeI restriction site in the SV40 PolyA that lacked the WPRE and cloned into the respective cut transgene cassette. Pfurin previously was described. Include pseudovirus plasmids.

### 10-1074 antibody production and purification

Briefly, HEK293T cells in T225 mm^2^ flasks were transfected with 60 ug total DNA/flask at a cellular 70% confluency with PEIpro transfection reagent. Cells were co-transfected with the AAV 10-1074 transfer plasmid and an additional plasmid encoding furin at a 4:1 ratio. At 16 hrs post-transfection, 10% FBS-DMEM media was replaced by serum-free 293 Freestyle media (Gibco). 48 hours following media replacement, media was collected, and debris was removed through centrifugation for 10 min at 1,500g and filtered using a 0.45 um flow filter unit (Thermo Fisher Scientific). Proteins were purified with HiTrap MabSelect SuRe columns (Cytiva) and eluted with IgG Elution Buffer (Thermo Fisher Scientific) into 1M Trizma HCL buffer, pH 9.0 (Millipore Sigma). Buffer was exchanged with 1X PBS and protein concentrated to 1-2 mg/mL with 30 kDa Amicon Ultra Centrifugation Filters (Millipore Sigma). Purified 10-1074 antibodies were assessed and confirmed by Coomassie-stained SDS-PAGE and stored at 4°C prior to use or -80°C for long-term storage.

### TZM-bl neutralization assay

TZM-bl neutralization assays were performed as previously described. Briefly, pseudovirus (BG505, PVO.4, CE1176, SIVmac239) was produced by transfecting HEK293T cells in T225 mm^2^ flasks at 70% cellular confluency. Cells were co-transfected with the respective pseudovirus envelope glycoprotein expression plasmid and the NL4.3ΔEnv expression plasmid at a 1:1 ratio using PEIpro. After an overnight incubation, medium was replaced with fresh 10% FBS-DMEM. Pseudovirus was harvested 48 hrs later and filtered using a 0.22 um syringe filter and stored at - 80°C until used. Pseudovirus was pre-incubated with titrated amounts of antibody in a 96-well plate with a starting concentration of 25 ug/mL in 10% FBS-DMEM for 1 hr at 37°C. TZM-bl cells were diluted in DMEM to 100,000 cells/mL and added to the virus/antibody mixture. Cells were then incubated for 44 hrs at 37°C. Viral entry was analyzed using Britelite Plus (Revvity) and luciferase was measured using a Synergy Neo 2 plate read (BioTek). For ex vivo neutralization analysis of macaque serum samples, heat inactivated serum samples were initially diluted 1:10, followed by five sequential 5-fold dilutions in 10% FBS-DMEM, and pre-incubated with pseudovirus (BG505, PV0.4, CE1176, SIVmac239) in a 96-well plate for 1 hour at 37°C. A negative control consisted of heat inactivated serum prior to AAV9 administration. TZM-bl cells were detached by trypsinization, diluted in DMEM to 100,000 cells/mL, and added to the virus/antibody mixture. Cells were then incubated for 44 hrs at 37°C. Viral entry was analyzed using Britelite Plus (Revvity) and luciferase was measured using a Synergy Neo 2 plate read (BioTek).

### AAV9 production and purification for mouse study

AAV9.10-1074 vectors were produced by triple transfection in HEK293T cells at 70% confluency. Expression plasmids encoding Rep2/Cap9, pHelper, and pAAV.10-1074 were mixed (1:1:1 ratio, 60 ug total) with PEIpro transfection reagent according to the manufacturer’s instructions. At 3-days post transfection, cells were harvested by adding 0.5M EDTA, pH 8.0 to each flask. Collected cells and medium were centrifuged for 10 minutes at 2,000g and washed with PBS. Cells were lysed using an AAV lysis buffer (150nM NaCl, 2mM MgCl2, 50mM Tris-HCL pH 8.0). Cells in lysis buffer were subject to three freeze/thaw cycles. Benzonase (50U/mL) and Triton X-100 (0.01%) were added to the lysate mixture and incubated for 1 hr at 37°C. The lysate mixture was then centrifuged for 1 hr at 7,000g and the supernatant was collected and filter sterilized using a 0.45um filter (Thermo Fisher Scientific). POROS Capture Select AAV9 columns (Cytiva) were used to purify AAV9 vectors. Buffer was exchanged with eluted vectors using 1X PBS and concentrated using 100 kDa Amicon Ultra Centrifuge Tubes. Vectors were quantified by qPCR using a Roche Lightcycler 480ii.

### AAV9 vector production for NHP study

AAV9.10-1074 vectors were produced at the University of Massachusetts Medical School Vector Core as previously described. Briefly, HEK293T were triple transfected with expression plasmids encoding Rep2/Cap9, pHelper, and pAAV.10-1074 encoding the respective promoter. After collecting cell lysates, AAV9 was purified through three sequential CsCl centrifugation steps. Vector genomes copy numbers (vg/mL) were determined through qPCR. AAV9 vectors were quality controlled by electron microscopy (EM) and the purity was verified through silver-stained SDS-PAGE.

### AAV vector administration in NSG mice

29 male NOD.Cg-Prkd^cscid^ IL2rg^tm1Wjl^/SzJ mice (NSG, strain number 005557) were obtained from The Jackson Laboratory. Mice were administered 2.5x10^12^ vg/kg of recombinant AAV9 vectors encoding 10-1074 at a 25 uL volume in the left gastrocnemius muscle. Mice were bled at weeks 1, 2, 3, 4, 6, and 8 post administration. About 20 uL of blood was collected, and plasma was obtained by centrifugation for 3 min at 11,000g. Plasma was frozen and kept at -80°C until used for analysis by gp120 ELISA. All mice were euthanized by at the end of the study in accordance to approved IACUC procedures.

### AAV vector administration in rhesus macaques

Prior to AAV9 administration, all macaques were prescreened for AAV9 neutralizing antibodies as previously described.^38^ Macaques were put into groups of four. Macaques were administered with a mixed dose of AAV9.10-1074 (1.25x10^12^ vg/kg), encoding the respective promoter, and AAV9.PD-L1 (1.25x10^12^ vg/kg) for a total dose of 2.5x10^12^ vg/kg. For the study evaluating 10-1074 expression without the WPRE, four macaques in this group were administered a mixed dose of AAV9.10-1074 (2.5x10^12^ vg/kg) and AAV9.PD-L1 (2.5x10^12^ vg/kg) for a total dose of 5x10^12^ vg/kg, matching the dose of the six macaques from our historical NHP study. Each macaque received eight i.m. injections: one in the lower left and right quadriceps muscle, one in the upper left and right quadriceps muscle, one in the left and right biceps muscle, and one in the left and right deltoids muscle. Blood draws were taken 2 weeks prior to AAV9 administration, weekly for the first four weeks, and bi-weekly up to 30 weeks post AAV9 administration. 11 macaques were released back to the colony at the conclusion of the study and 5 macaques were euthanized for sample harvesting in accordance to approved IACUC procedures.

### 10-1074 gp120 ELISA

Half-area 96-well Costar Assay Plates (Corning) were coated with 5 ug/mL of ADA gp120 (Immune Tech) in PBS and kept overnight at 4°C. Following overnight incubation, plates were washed twice with PBS-T (PBS + 0.05% Tween-20) and blocked with blocking buffer (PBS + 5% BSA + 5% milk + 0.1% Tween 20) for one 1 hr at 37°C. Plasma samples from mice or serum from macaques were unblinded for gp120 ELISA. Samples were serially diluted in blocking buffer and were added to the plate in duplicate. Standard curves were generated by diluting 4 ug/mL of recombinant 10-1074 in blocking buffer and plated in duplicate. Plates were incubated for 1 hr at 37°C. Plates were washed five time with PBS-T and a horseradish peroxidase secondary antibody targeting the IgG Fc (Jackson ImmunoResearch). Plates were incubated for 1 hr at 37°C and then washed ten times with PBS-T. TMB solution (Thermo Fisher Scientific) was added and plates were incubated at room temperature until their standard curves developed (typically 2-3 minutes). TMB Stop Solution (SeraCare) was added, and absorbance was read at 450 nm by a Synergy Neo2 multi-mode reader (BioTek). Standard curves were analyzed using GraphPad Prism 10.5.0 software and used to determine protein titers from plasma and sera samples.

### Anti-drug antibody ELISA

Macaque sera samples were unblinded for ADA ELISAs. Sera samples from macaques expressing 10-1074 were analyzed for anti-10-1074 antibodies. 96 half-well plates were coated with 5 ug/mL of recombinant 10-1074 in PBS and incubated overnight at 4°C. Following overnight incubation, plates were washed twice with PBS-T and blocked with blocking buffer for 1 hr at 37°C. Sera samples were diluted 1:5 in blocking buffer, diluted 5-fold to a 1:1250 dilution, and plated in duplicate for 1 hr at 37°C. Plates were then washed five times with PBS-T. Anti-10-1074 antibodies were measured using a secondary antibody detecting the kappa light chain (Millipore) at a 1:5000 dilution. Plates were incubated for 1 hr at 37°C and washed ten times with PBS-T. TMB solution was added and plates were incubated at room temperature for 4-5 minutes. TMB Stop Solution was added and absorbance at 450 nm was read as described above.

### IFN-γ ELISpot assay

96-well HA 0.45μm Multiscreen Filter Plates (Millipore) were coated with 5 μg/mL of mouse anti-human IFN-γ and incubated overnight at 4°C. Following overnight incubation, plates were washed four times and blocked with complete lymphocyte medium (RPMI 1640 w/ L-glutamine + 10% FBS + 10% HEPES buffer + 1% penicillin-streptomycin) for 1h at 37°C. Prior to 1 hr block, isolated PBMCs were thawed and washed with lymphocyte medium with Benzonase (50 U/mL). After Benzonase treatment, PBMCs were resuspended in lymphocyte medium and incubated for 2 hrs at 37°C. Previously described peptide pools (PD-L1, LS-Furin P2A, 10-1074 VarL, 10-1074 VarH) were diluted in lymphocyte medium to a final concentration of 10 μg/mL. Staphylococcal enterotoxin B (ToxTech) was used a as positive control at 2 μg/mL and DMSO (Sigma) was used as negative control. After plating peptides, 200,000-250,000 PBMCs were added to wells in duplicate and incubated for 20 hrs at 37°C. Following incubation, plates were washed 4 times PBS-T and incubated with 1 μg/mL Biotin-α-human IFN-γ mAb diluted in PBS-T + 1% FBS (PBST-FBS) for three hours at 37°C. Plates were then washed four times with PBS-T and incubated with streptadavin-conjugated alkaline phosphatase (Rockland) (1:1000 dilution in PBST-FBS) for 1 hr at 37°C. Lastly, plates were washed four times with PBS-T and One-Step NBT/BCIP (Thermo Fisher Scientific) was added to develop IFN-γ spots. Analysis was performed using the CTL ImmunoSpot analyzer and single-color suite software (Ver. 7.0; Cellular Technology Ltd).

### AAV neutralization assay

Assays were performed as previously described. Briefly, heat inactivated macaque serum samples were diluted 1:5, followed by five sequential 4-fold dilutions in 10% FBS-DMEM and added to a 96-well plate in duplicate. AAV.CAG.fLuc vector was added and diluted in 10% FBS-DMEM to a concentration (5x10^9^ to 2x10^11^ vg/mL depending on AAV capsid used) that would result in RLUs greater than 10,000. AAV vector was then added to the 96-well plate with serum and incubated for 1 hr at 37°C. HEK293T cells were diluted in DMEM to 30,000 cells/mL and added to the serum/AAV mixture. Cells were then incubated for 24 hrs at 37°C. AAV neutralization was analyzed using Britelite Plus (Revvity) and luciferase was measured using a Synergy Neo 2 plate read (BioTek).

### Statistical Analysis

ANOVA (one-way & two-way), Tukey’s multiple comparison test, unpaired t-test, and correlation analysis were performed using GraphPad Prism v10.6.0. P values <0.05 were considered statistically significant. Pearson r values between 0.5 and 0.7 were moderately correlated and values >7.0 were highly correlated.

## Acknowledgements

The authors would like to thank the Emory National Primate Research Center (ENPRC) staff for their tremendous help in completing this study; and M.E. Davis-Gardner for her comments and edits to the manuscript. This work was supported in part by National Institutes of Health awards R01AI167724 (M.R.G.) and R01DA056770 (M.R.G.). Additional support was provided from the NIH Office of Research Infrastructure Programs (ORIP) P51OD11132 to ENPRC and U42OD011023 to ENPRC.

## Author Contributions

I.L, M.R.G designed the study. I.L, A.A.K, and N.S.C performed both in vitro and in vivo experiments. I.L, A.A.K, P.D, and Y.B were responsible for blood processing in both mice and NHP studies. M.K provided critical reagents and provided input into experiments performed for the NHP study. I.L conducted all data analysis and prepared the initial draft of the manuscript. All co-authors provided feedback, contributed to manuscript revisions, and approved the submitted manuscript.

## Competing Interests

IL, AAK, MK, and MRG are named inventors on a patent application related to the technologies described in this study submitted by Emory University. MRG is a co-founder and consultant for Emmune, Inc. MRG has consulted for ViiV Healthcare. GG is a co-founder of Voyager Therapeutics and Aspa Therapeutics and holds equity in both companies. GG is an inventor on patents with potential royalties licensed to Voyager Therapeutics, Aspa Therapeutics, and other biopharmaceutical companies. The remaining authors declare no competing interests regarding this study.

---

**Figure S1.**
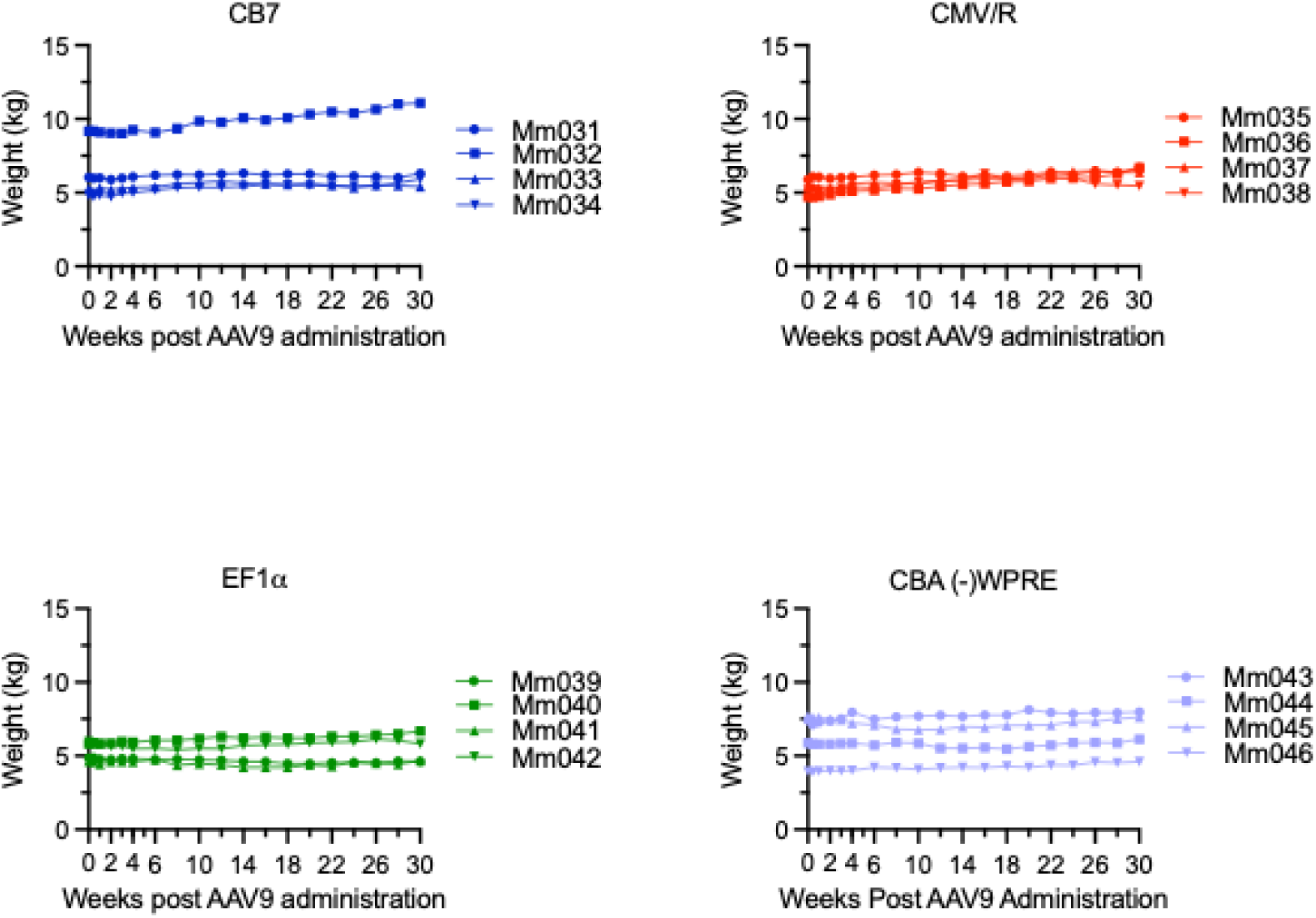
Rhesus macaque weights following co-delivery of AAV9.10-1074 and AAV9.PD-L1. Body weight (kg) as determined for each indicated time point post AAV9 administration. No sustained or clinically significant weight loss was observed in any group.

**Figure S2.**
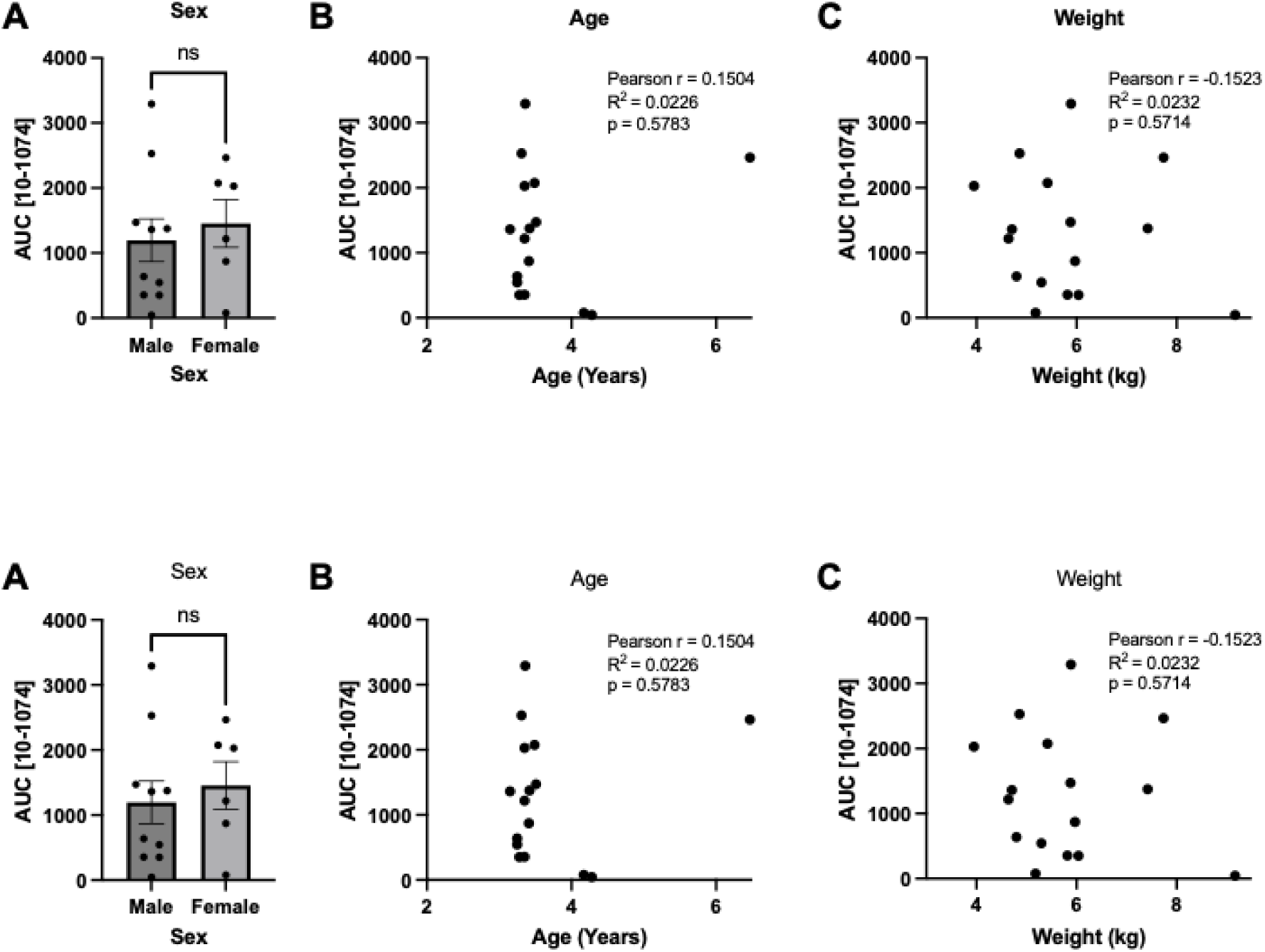
Correlation analysis of 10-1074 concentration with sex, age, and weight in rhesus macaques. Area under the curve (AUC) of serum 10-1074 concentrations in all 16 macaques was compared across sex (**A**), age (**B**), and weight (**C**) following co-delivery of AAV9.10-1074 and AAV9.PD-L1. There were no significant associations observed for any comparison. Statistical analysis for (A) was determined by an unpaired two-tailed t test with each dot representing the AUC for each animal. n.s. indicates not significant. Statistical analysis for (B) and (C) was determined by a Correlation test. Pearson r values, p-value (two-tailed), and R^2^ values are included.

**Figure S3.**
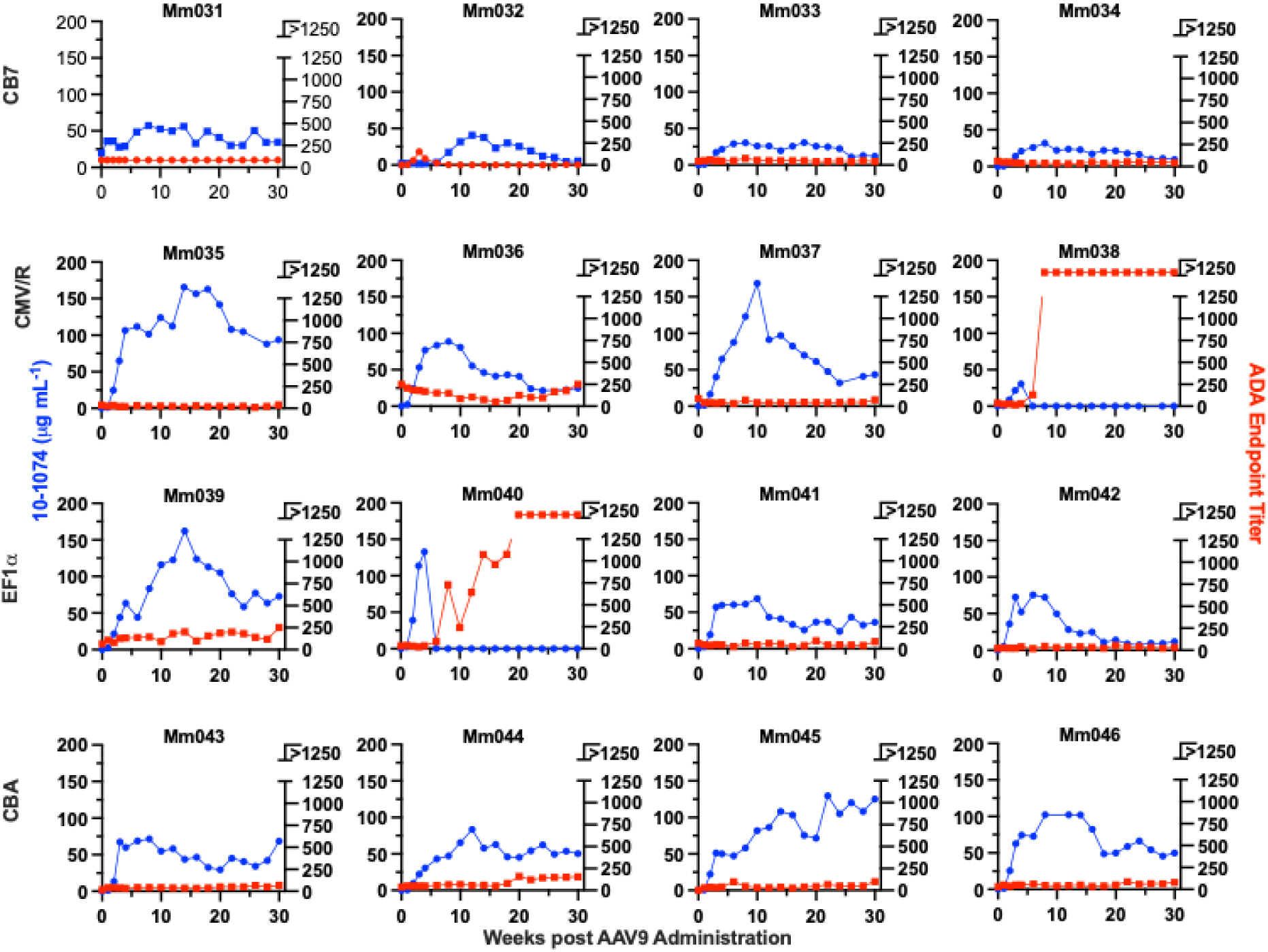
Individual rhesus macaque 10-1074 serum concentrations and ADA endpoint titers. 10-1074 serum concentrations (blue) and ADA (red) against 10-1074 for each macaque throughout the 30-week study. Promoter groups are labeled for each row.

**Figure S4.**
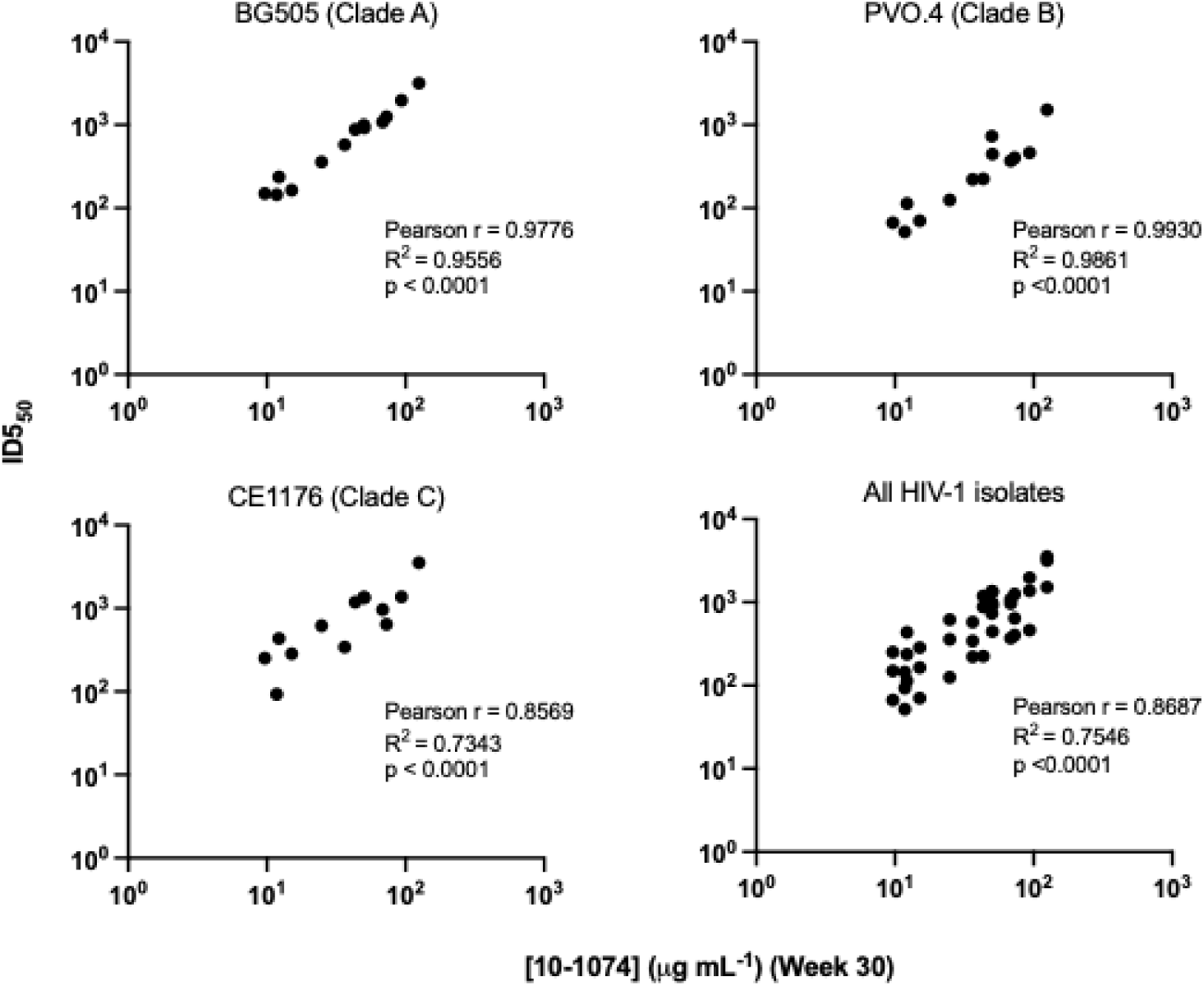
Correlation between 10-1074 serum concentration 30 weeks post-AAV9 administration and neutralization potency. Correlation analyses were performed between neutralization potency (obtained from ex-vivo neutralization assays) and measurable week 30 serum 10-1074 serum concentrations of all rhesus macaques. Each dot represents an individual macaque. ID_50_ values represent the reciprocal serum dilution achieving 50% inhibition of infection. Statistical analysis was determined by a Correlation test. Pearson r values, p-value (two-tailed), and R^2^ values are included.

